# The influence of relationship closeness on default-mode network connectivity during social interactions

**DOI:** 10.1101/815274

**Authors:** Dominic S. Fareri, David V. Smith, Mauricio R. Delgado

## Abstract

Reciprocated trust plays a critical role in forming and maintaining relationships, and has consistently been shown to implicate neural circuits involved in reward-related processing and social cognition. Less is known about neural network connectivity during social interactions involving trust, however, particularly as a function of closeness between an investor and a trustee. We examined network reactivity and connectivity in participants who played an economic trust game with close friends, strangers and a computer. Network reactivity analyses showed enhanced activation of the DMN to social relative to non-social outcomes. A novel network psychophysiological interaction analysis (nPPI) revealed enhanced connectivity between the DMN and the superior frontal gyrus and superior parietal lobule when experiencing reciprocated vs. violated trust from friends relative to strangers. Such connectivity tracked with differences in self-reported social closeness with these partners. Interestingly, reactivity of the executive control network (ECN), involved in decision processes, demonstrated no social vs. non-social preference, and ECN-ventral striatum (VS) connectivity did not track social closeness. Taken together, these novel findings suggest that DMN interacts with components of attention and control networks to signal the relative importance of positive experiences with close others vs. strangers.

## Introduction

Trust and reciprocity are cornerstones of forming and maintaining close relationships (reviewed in Simpson, 2007; Krueger and Meyer-Lindenberg, 2019). Indeed, deciding whether to trust or reciprocate generosity depends upon our ability to integrate prior experiences and knowledge about others with future expectations of them (reviewed in Fareri, 2019). Substantial evidence suggests that regions supporting social cognition and reward circuitry (e.g., medial prefrontal cortex (mPFC), striatum) help to distinguish between close and distant others and integrate prior expectations with future choices during social interactions (Krienen *et al.*, 2010; Fareri *et al.*, 2012; Tamir and Mitchell, 2012; Fareri *et al.*, 2015). However, the way in which close relationships shape the function and connectivity of large-scale networks during social interactions is largely unknown, which is a critical point to consider given recent work demonstrating representation of social space in a network of ‘social brain’ regions (Parkinson *et al.*, 2017; Parkinson *et al.*, 2018).

We have previously demonstrated that information about relationship closeness can shape the representation of shared positive experiences and of reciprocity at the behavioral and neural levels. The ventral striatum shows enhanced activation when earning a shared monetary reward with a close friend relative to a stranger (Fareri, Niznikiewicz, *et al.*, 2012), and people compute an added social value when experiencing reciprocity from a friend contingent upon interpersonal aspects of the friendship (Fareri *et al.*, 2015). These findings have built upon extant literature suggesting that neural circuits supporting reward valuation (e.g., striatum, ventral mPFC; Haber and Knutson, 2010; Bartra *et al.*, 2013) are also sensitive to social information (for review see Fareri and Delgado, 2014; Ruff and Fehr, 2014), representing the value of experienced and anticipated social rewards (e.g., social approval, praise) (Izuma *et al.*, 2008; Mobbs *et al.*, 2009; Izuma *et al.*, 2010; Smith *et al.*, 2010) and the encoding of probabilistic social feedback (i.e., acceptance, rejection) (e.g., Somerville *et al.*, 2006; Jones *et al.*, 2011).

While relationships clearly shape neurocomputational signals of social value, much less is known about how the context of close relationships influences the response and connectivity of *networks* of regions – in particular, networks that respond to social information during trust-based interactions. The DMN, which is often most active at rest (Raichle *et al.*, 2001; Buckner and Vincent, 2007; Buckner and DiNicola, 2019), is comprised of structures including the precuneus, mPFC, posterior cingulate cortex (PCC) and temporoparietal junction. While the DMN is posited to be involved in self-referential thought, it has also been theorized to generate predictions based on past experiences to inform future action (Barrett, 2017), and its subcomponents are all frequently implicated in social cognition (Adolphs, 2009; Stanley and Adolphs, 2013). Together, these findings have prompted additional hypotheses that a major function of the default state of the brain is to prime us for social cognition (Schilbach *et al.*, 2008; Spunt *et al.*, 2015; Meyer, 2019). Yet, how and to where such social information gets communicated *during* social interactions is not well-established.

A host of networks may be involved in representing and integrating information about close others during social interactions (Laurita *et al.*, 2019). One possibility is that the DMN interacts with reward circuitry to integrate the context of a close relationship with experienced reciprocity in computing social reward value, as both activation in both mPFC and ventral striatum correlate with computational social value signals (e.g., Fareri *et al.*, 2015). An alternative possibility is that the DMN may interact with components of control and attention-related networks (e.g., executive control network (ECN), salience network, frontoparietal network) to integrate information about the context of close relationships and inferred intentions of close others (relative to strangers) during trust-based interactions (Igelström *et al.*, 2016; Monfardini *et al.*, 2016; Bellucci *et al.*, 2019). The DMN may thus draw on past experiences to aid in social prediction (Barrett, 2017). Relatedly, other networks, such as the ECN—which includes lateral PFC, anterior insula, mPFC (anterior and paracingulate cortices; Smith *et al.*, 2009) and is implicated in goal-directed behavior more generally—may be involved with processing positive and negative outcomes more generally. This would be consistent with the idea that social processes may involve sub-networks that may be exclusive from those supporting more general cognitive function (Meyer and Lieberman, 2012).

The goal of this study was to characterize how close relationships influence network function and connectivity during trust-based social interactions with friends and strangers. We employed a novel form of psychophysiological interaction analysis (Friston *et al.*, 1997)—network PPI (nPPI) (Utevsky *et al.*, 2017). Unlike seed-based approaches, which examine connectivity between specific voxels or a specific region and the rest of the brain, nPPI capitalizes on the dynamics of entire neural networks and allows for the examination of network connectivity during task conditions. We hypothesized that the DMN would demonstrate increased reactivity to social relative to non-social (i.e., computer) conditions, and that this might be heightened as a function of the relationship one has with a partner (i.e., friend > stranger). Based on our prior work, we further hypothesized that the DMN would exhibit increased connectivity with reward-related circuitry (i.e., ventral striatum) when experiencing reciprocity from a close friend relative to a stranger, and that this pattern would vary with the degree of social closeness felt towards friends and strangers. We also tested whether the executive control network (ECN), which has been shown to facilitate goal-directed processes in general as well as orientation towards socially relevant stimuli (Seeley *et al.*, 2007; Smith *et al.*, 2009; Utevsky *et al.*, 2017) demonstrated differential reactivity and connectivity during social interactions for friends relative to strangers. However, we did not expect that this would vary with self-reported social closeness.

## Methods

### Participants

Secondary data analyses were conducted on data collected from 26 participants (14 F/12 M, mean age = 21.36, sd = 3.67) in a previously published study from our group (Fareri *et al.*, 2015). Participants had no history of head trauma or psychiatric illness and all participants provided informed consent. All procedures were approved by the Rutgers University Institutional Review Board.

### Experimental Paradigm

All participants played an iterated economic trust game with three different partners—a same-sex close friend, a same-sex stranger (laboratory confederate) and a computer (Fareri et al., 2015). Participants were asked to bring a same-sex close friend with them to the experimental session at the Rutgers University Brain Imaging Center (RUBIC, Newark, NJ). Participants and their friends met a same-sex confederate (stranger) at the imaging center who was portrayed as another participant in the study, but was actually a member of the laboratory team. Briefly, MRI participants were in the role of investor in the trust game (Berg et al., 1995; King-Casas et al., 2005) and the three partners were in the role of trustee. Participants chose whether to invest money with or keep money from one of their three partners on each trial (event-related design, partner presentation was randomly presented). After submitting their decision, if they chose to invest with a partner, participants experienced a jittered ISI during which they awaited their partner’s response (i.e., reciprocity, violation of trust). All participants were trained together on the trust game task in the scanner control room, and were under the impression they would be interacting in real time over networked computers between the scanner and control room. In reality, friends’ and strangers’ behavior was preprogrammed into the task script (E-Prime 2.0; Psychology Software Tools, Pittsburgh, PA), such that all partners provided 50% reinforcement rates (i.e., partners reciprocate and defect with equivalent probability) on trials in which the MRI participant decided to invest (see Fareri et al., 2015 for complete details). Prior to the start of the task, MRI participants filled out the Inclusion of Other in Self Scale (IOS), a self-report questionnaire assessing social closeness (Aron et al., 1992) with their three partners (friend, stranger, computer).

### fMRI Acquisition and Preprocessing

Neuroimaging data were acquired on a 3T Siemens Magnetom Trio whole-body scanner at the Rutgers University Brain Imaging Center (RUBIC; Newark, NJ). Structural images were collected using a standard T1-weighted MPRAGE sequence (256 × 256 matrix; FOV = 256mm; 176 1mm sagittal slices). Functional images were acquired using a single shot gradient EPI sequence (TR = 2000ms, TE = 30ms, FOV = 192, flip angle = 90°, bandwith = 2232 Hz/Px, echo spacing = 0.51), comprising 33 oblique-axial slices (3 × 3 × 3mm voxels) parallel to the anterior-posterior commissure line, collected in an ascending-interleaved order. Data were preprocessed using a combination of custom scripts (https://github.com/rordenlab/spmScripts) for SPM12 and FSL (v5.09; FMRIB). Standard preprocessing steps including motion correction, brain extraction and coregistration were performed in SPM. Functional data were acquired in an ascending-interleaved fashion; slice-time correction was performed in SPM aligning to the first slice. Motion artifact was addressed through an automated independent components analysis approach employing ICA-AROMA in FSL (Pruim *et al.*, 2015). This approach identifies noisy components in single subject functional data by computing the degree to which each component is characterized by characteristic patterns of motion artifact (i.e., high frequency signals, correlation with standard realignment parameters, overlap with CSF and edge voxels); components that highly identify with these characteristic patterns are removed from single subject functional data at the single run level via linear regression. The resulting denoised data serves as input to first level models. We additionally computed estimates of frame-to-frame motion using MCFLIRT in FSL to compute relative mean framewise displacement, which were subsequently used as an additional group level covariate in offline analyses of relationships between connectivity and behavior (see below); this allowed for assessment of whether observed results were not due solely to motion.

### Network Psychophysiological Interaction Analysis (nPPI)

We employed a novel connectivity approach, nPPI, (Utevsky *et al.*, 2017) aimed at assessing changes in task-based connectivity of canonical neural networks. nPPI improves upon seed-based connectivity approaches by leveraging the fact that the brain is organized into functional neural networks during both task and rest states (Smith *et al.*, 2009). nPPI treats entire functional networks as ‘seeds’ to capture functional network dynamics and maps those to interactions with other regions of the brain at voxel level. Specifically, we captured network dynamics using a spatial regression, where the functional data were regressed onto a 4-dimensional design matrix consisting of the 10 canonical networks from prior work (Smith *et al.*, 2009). Notably, this process is identical to the first stage of the popular dual-regression analysis (Filippini *et al.*, 2009; Nickerson *et al.*, 2017) and extends it by including psychophysiological interactions with specific networks of interest (Friston *et al.*, 1997; Smith *et al.*, 2016) while controlling for the dynamics of other networks in the analysis (McLaren *et al.*, 2012).

We were interested in investigating whether the default mode network (DMN) demonstrates differential patterns of effective connectivity during social interactions involving close friends and strangers. Specifically, we focused on how DMN connectivity was altered during the processing of positive (i.e., reciprocity) vs. negative (i.e., defection) trust game outcomes as experienced from these different partners. We extracted time series from 10 canonical resting state networks as identified by Smith and colleagues (Smith *et al.*, 2009). We constructed a GLM using FEAT with regressors at the first level that modeled: the decision phase (agnostic to partner and participant choice), and outcome phase (separate regressors for experienced reciprocity and defection from each partner), and nuisance regressors modeling missed trials and the outcome phase of trials in which participants defected. The first-level nPPI models also included the physiological timeseries from each network (10 total). We constructed separate first-level generalized nPPI models to examine connectivity of the DMN and the ECN, such that the PPI regressors in one model were comprised of interactions between the DMN timeseries and task regressors, while in the other we modeled interactions between the ECN timeseries and task regressors. First level GLMs were conducted at the single run level, and were combined for each subject at the second level. Group-level whole-brain analyses were corrected for multiple comparisons using permutation testing (10000 permutations) and Threshold-Free Cluster Enhancement as implemented in FSL with variance smoothing (set to 2.13) via randomise. Group-level analyses included a mean-centered subject-level covariate representing the difference in self-reported social closeness between MRI participants and their friends relative to strangers (i.e., IOS Friend - IOS Stranger).

To initially characterize the profile of the functional response of canonical networks—DMN, ECN—during the outcome phase of the task, we regressed the timeseries of each of these networks on a simpler model including only the 10 task-based regressors (Decision Phase: Keep, Share; Outcome Phase: Reciprocate, Defect for each partner, participant Keep, Missed Trials). Parameter estimates indexing the response of each network during trust game outcomes with each partner were extracted and plotted, with differences between partner and outcome conditions tested using a repeated measures ANOVA in jamovi (jamovi.org).

## Results

### Enhanced DMN reactivity for social outcomes

In order to establish whether the DMN may be preferentially sensitive to social relative to non-social outcomes in comparison to the ECN, we first examined the reactivity of these networks during the outcome phase of the trust game. Regression of network timeseries from the DMN and ECN revealed divergent patterns of reactivity (see Figure 2). A 2 (Network) × 3 (Partner) × 2 (Outcome) repeated measures ANOVA revealed a significant Partner x Network interaction (*F*_(2,50)_ = 11.88, p < .001, Mauchly’s test of Sphericity: W = .996, p = 0.95). Participants demonstrated stronger recruitment of the DMN compared to the ECN when experiencing outcomes with a close friend (*t*_(58.2)_ = 4.38, p<.001) and a stranger (*t*_(58.2)_ = 4.38, p < .001), but not when experiencing outcomes with a computer (*t*_(58.2)_ = −0.32, p > 0.75). This analysis also revealed a significant Outcome x Network interaction (*F*_(1,25)_ = 5.14, p < 0.04). Participants demonstrated stronger activation of the DMN compared to the ECN when experiencing reciprocity (*t*_(37.9)_ = 4.30, p < .001), but not violations (*t*_(37.9)_= 2.19, p = .104), of trust. Post-hoc tests were corrected using a sequential Bonferroni correction (Holm, 1979). All other interactions were non-significant (p’s > 0.35). We note that the finding of enhanced DMN reactivity to social relative to non-social outcomes holds when examining the DMN in isolation using a 2 (outcome) × 3 (partner) repeated measures ANOVA (Main Effect of Partner: F _(2,50)_ = 34.36, p<.001, Mauchly’s test of Sphericity: W = .951, p = .549; Friend > Computer, p <.001; Stranger > Computer, p <.001).

**Figure 1.**
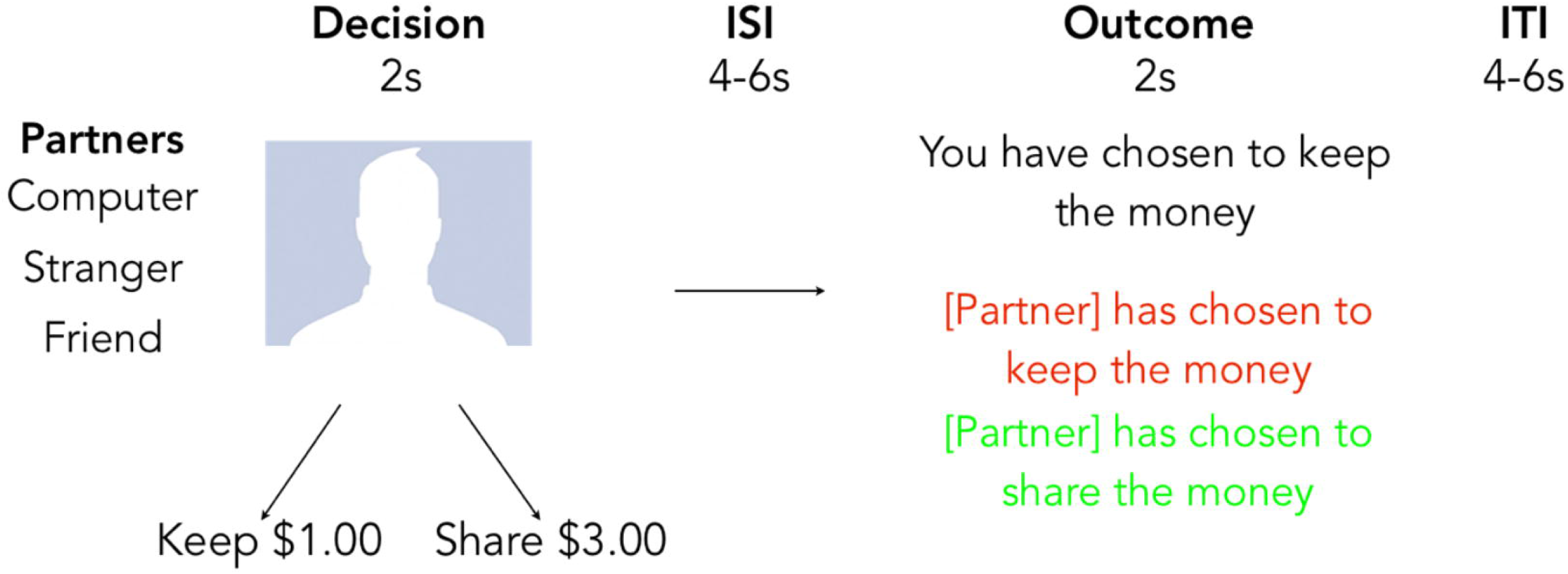
Task Schematic. Participants played an iterated economic trust game with 3 different partners: a computer, a same-sex stranger, and a same-sex close friend (see Fareri*, Chang* and Delgado, 2015).

**Figure 2.**
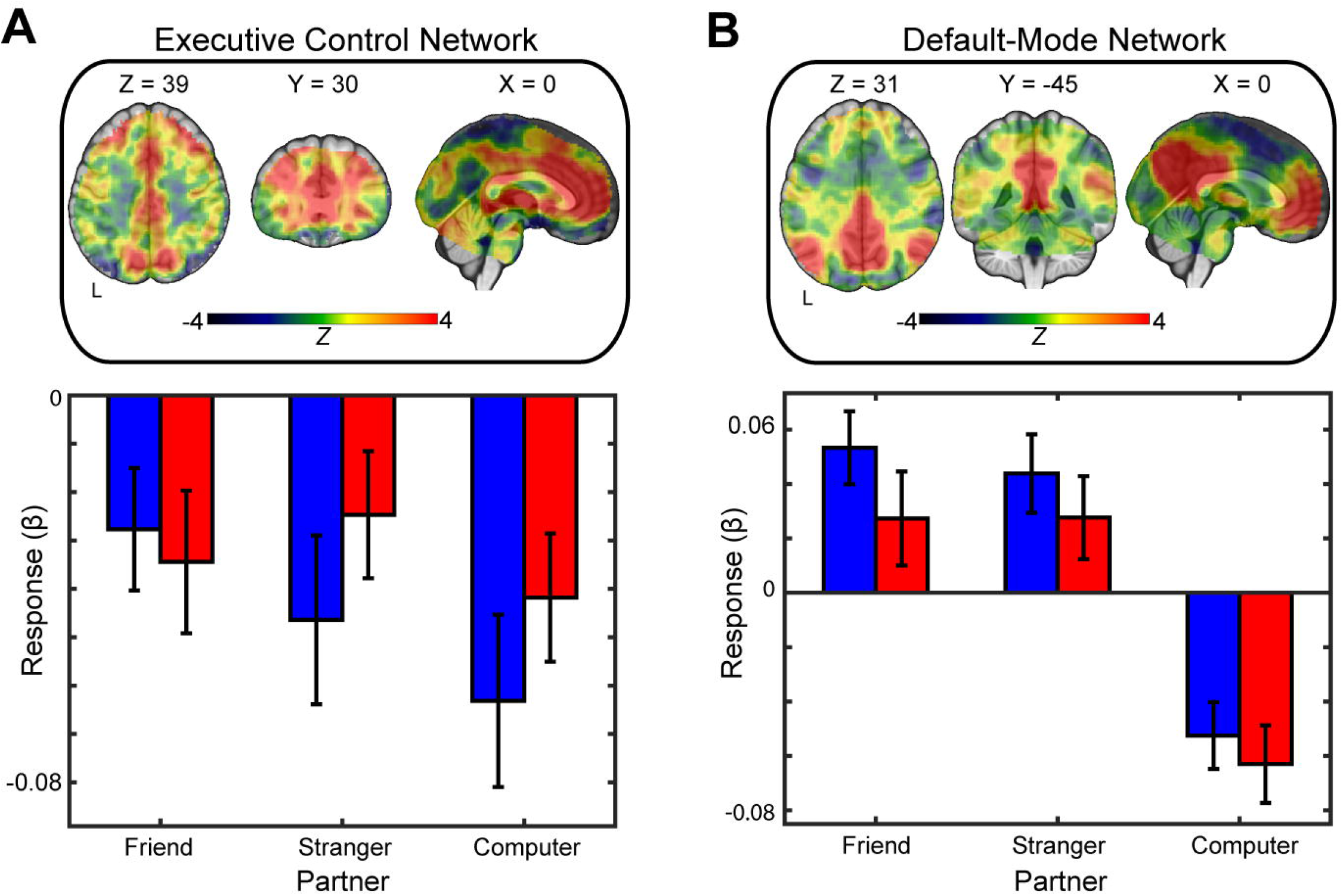
Network Reactivity. Regression of DMN and ECN timeseries on task conditions during social outcome processing revealed enhanced reactivity for social relative to non-social outcomes in the DMN, but not the ECN (Green bars = reciprocation of trust; Red bars = violation of trust). Warm colors on the brain maps represent regions that comprise the networks of interest.

As an additional post-hoc analysis, we tested whether the DMN timecourse was being driven by a single node of the network. We extracted the raw timecourses (i.e., task-independent) from peak seed locations within each of the mPFC (2,56,−4), bilateral TPJ (left: −44, −60, 24; right: 54, −62, 28) and PCC/Precuneus (2, −58, 30). Results indicated that the timecourse of each node of the network was highly correlated with the timecourse of the entire DMN, with the PCC/Precuneus being the most strongly correlated (mPFC: r = .62; lTPJ: r = .70; rTPJ: r = .69; PCC/Precuneus: r = .84; see Supplemental Figure 1a). After partialling out the contributions of other nodes within the network, however, we note that PCC/Precuneus appeared to contributing the most to the DMN timecouse (mPFC: r = .25; lTPJ: r = .32; rTPJ: r = .17; see Supplemental Figure 1b; Utevsky *et al.*, 2014).

### DMN connectivity during social outcomes is sensitive to relative differences in social closeness

Based on both our prior work (Fareri *et al.*, 2012; Fareri *et al.*, 2015) and the above results demonstrating increased DMN reactivity to social relative to non-social trust game outcomes, we next investigated whether network connectivity during social interactions changed as a function of both partner and social outcome valence. We conducted a network PPI (nPPI) analysis to address this question, though we note that we interpret these results cautiously given that our sample size is less than the minimum typically suggested (n=40) to assess brain-behavior relationships (Yarkoni and Braver, 2010). Specifically, we aimed to investigate whether the DMN would show *increased* connectivity when experiencing social reward (reciprocity) compared to social punishment (defection) from friends relative to strangers. We tested this question by conducting a double subtraction of (Friend Reciprocate > Friend Defect) > (Stranger Reciprocate > Stranger Defect). Importantly, given our prior findings showing that between subject differences in social closeness with friends and strangers plays a role in neural representations of social outcomes (Fareri *et al.*, 2012; Fareri *et al.*, 2015), we also included a between-subjects social closeness covariate (Friend – Stranger social closeness difference score, mean centered) for each participant. This analysis revealed significant coupling of the DMN with the superior frontal gyrus (x, y, z (MNI) = 18, 46, 40) and superior parietal lobule (x, y, z (MNI) = 10, 22, 42) (p<.025, whole-brain corrected with TFCE; Figure 3). For visualization purposes, we extracted connectivity estimates from these clusters and plotted them against the difference in reported social closeness between friends and strangers; scatterplots (Figure 3) demonstrate that greater closeness exhibited towards a friend (vs. a stranger) was associated with more positive coupling between the DMN and these regions. Similar patterns emerged for clusters in the fusiform gyrus (x, y, z = 17, 12, 21) and the lingual gyrus (x, y, z = 32, 19, 19), but are not depicted here. Interestingly, conducting the same double subtraction using the ECN as a seed network revealed no clusters representing significant connectivity.

**Figure 3.**
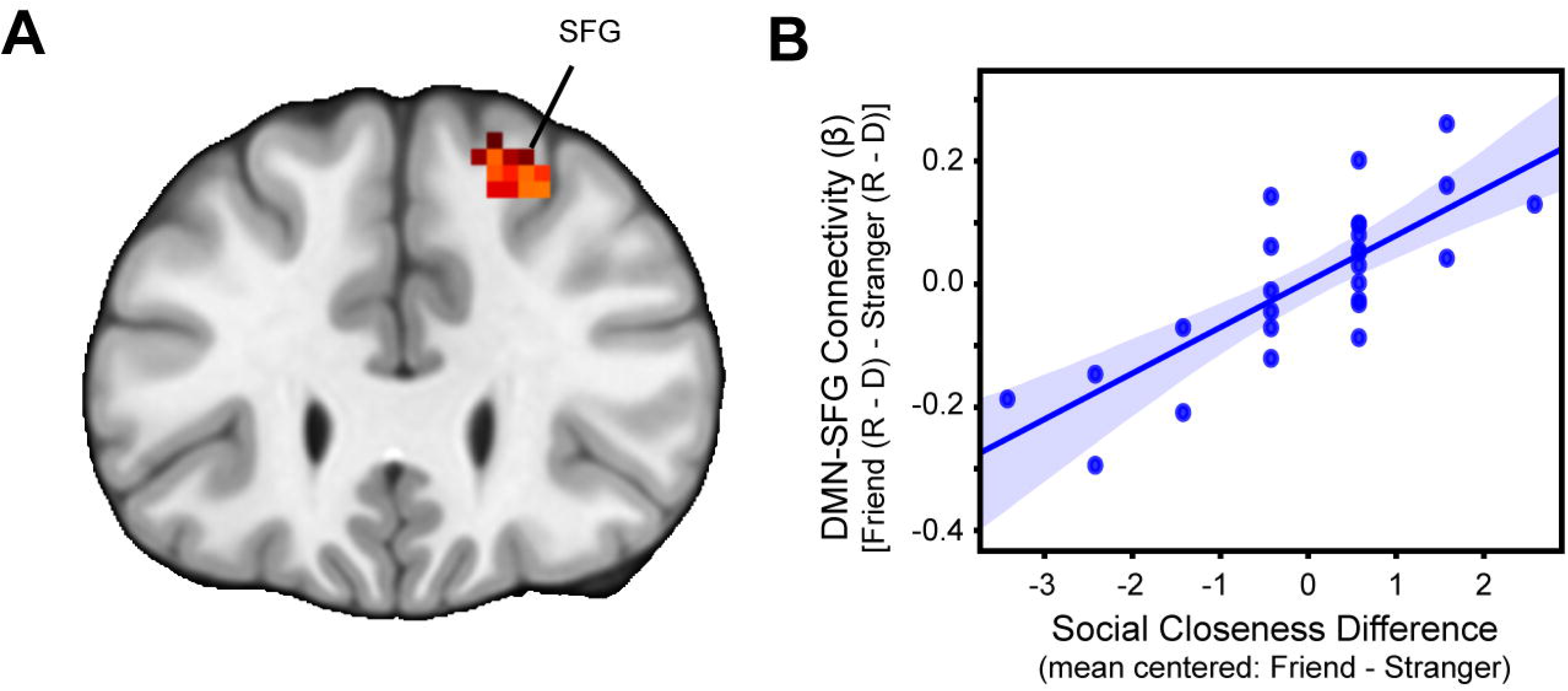
nPPI: DMN Connectivity. Network PPI revealed enhanced coupling of the DMN with superior frontal gyrus when participants experienced reciprocity relative to violations of trust from a friend relative to a stranger, which correlated with differences in self-reported social closeness (IOS) with friends vs. strangers.

### Connectivity with reward circuitry during interactions with a friend

Given our prior work demonstrating a link between activation in mPFC and the ventral striatum during experienced reciprocity from a friend (Fareri *et al.*, 2015), we conducted targeted PPI contrasts examining both DMN and ECN connectivity during reciprocity (vs. defection) experienced with a close friend specifically. Interestingly, these analyses revealed no voxels exhibiting significant connectivity with the DMN. However, we did find enhanced connectivity between the ECN and bilateral striatum when experiencing reciprocity from a close friend relative to a stranger at a slightly more lenient threshold (TFCE, p<.05; see Figure 4). As an exploratory measure, we also extracted connectivity estimates from these same ventral striatal voxels (x, y, z = 3, 11, −4) during experiences with a stranger; a post-hoc 2 (outcome) × 2 (partner) repeated measures ANOVA revealed no difference in ECN-VS connectivity during experiences with a friend relative to a stranger (*F*_(1,25)_ = 0.02, p > 0.9) and no interaction (*F*_(1,25)_ = 1.69, p<0.2).

**Figure 4.**
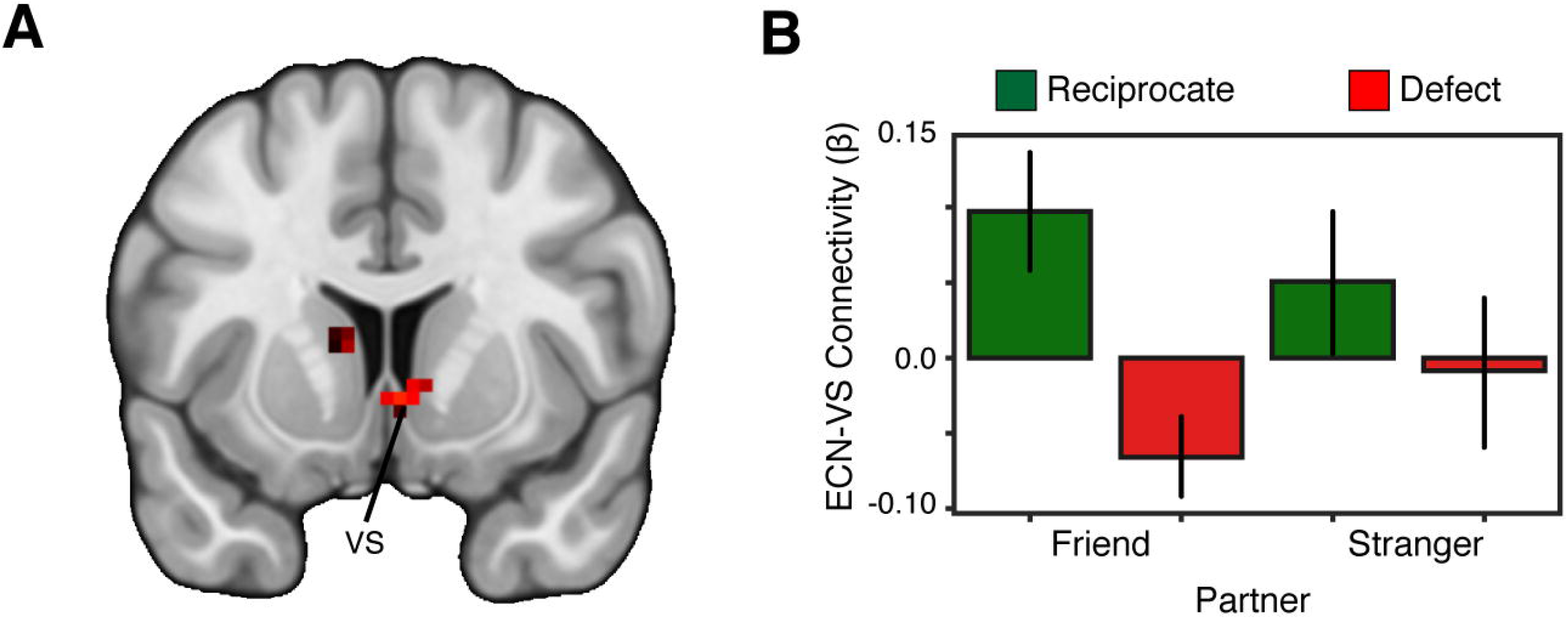
nPPI: ECN Connectivity. Targeted nPPI analyses during interactions with a close friend using the ECN as a seed region revealed enhanced coupling with the bilateral striatum during reciprocity relative to violations of trust. Post-hoc tests indicated no significant difference in ECN connectivity with the ventral striatum during interactions with a stranger.

## Discussion

The goal of this study was to investigate how the closeness of a social relationship shapes the connectivity of canonical neural networks during experiences of reciprocity and violations of trust. Taking a network-based approach, our findings demonstrated a preference for social relative to non-social outcomes in terms of the DMN response to experienced reciprocity relative to the ECN. Novel nPPI analyses further revealed that the DMN exhibits enhanced connectivity with later prefrontal and lateral parietal regions when experiencing reciprocity relative to violations of trust from a friend relative to a stranger, which is more positive the closer one feels to a friend. Consistent with network reactivity analyses, the ECN did not demonstrate differential connectivity as a function of outcome valence and relationship context. Interestingly, targeted analyses revealed that the ECN did show significant connectivity with the striatum bilaterally, specifically when experiencing reciprocity relative to violations of trust from a friend only, whereas this pattern of connectivity with the striatum was not observed in the DMN. Taken together, our results suggest that: 1) the DMN demonstrates a preference for socially relevant outcomes; and that 2) it may interact with components of networks involved in attention and cognitive control to differentially represent the importance of social experiences based on relationship closeness.

The role of the DMN has been classically viewed through the lens of resting-state activation and connectivity. Given that this network demonstrates robust activation at rest in task-negative states (Raichle *et al.*, 2001; Utevsky *et al.*, 2014), it has been suggested that the DMN plays a significant role in self-referential processing (Gusnard *et al.*, 2001), memory consolidation and maintenance of intrinsic neural relationships (Buckner and Vincent, 2007). However, an alternative hypothesis built around findings that the DMN shows increased activation during self and social processes (reviewed in Buckner and DiNicola, 2019) suggests that the DMN may prime us for social function, and that social cognition may in fact be our brain’s default state (Meyer, 2019). Regions comprising the DMN (i.e., mPFC, PCC/precuneus, TPJ) are broadly implicated in self-other representation, mentalizing/theory of mind, social perception, social appraisals and decision-making (Adolphs *et al.*, 1998; Amodio and Frith, 2006; Mitchell *et al.*, 2006; Adolphs, 2010; Krienen *et al.*, 2010; Koster-Hale and Saxe, 2013; Stanley and Adolphs, 2013; Deen *et al.*, 2015; Spunt *et al.*, 2015; Stanley, 2016). Activation within the DMN during rest components of a task (i.e., fixation) and when engaged in mentalizing show significant overlap (Schilbach *et al.*, 2012; Spunt *et al.*, 2015) and are posited to form a social-affective sub-network implicated in social cognition and emotional processes (Amft *et al.*, 2015; Eickhoff *et al.*, 2016; Alcalá-López *et al.*, 2018). Our finding that the DMN shows increased task-based reactivity (relative to the ECN) when processing social versus non-social outcomes supports the idea that the DMN preferentially engages in social cognitive processes involving self-other representation. Further, the increased reactivity of the DMN relative to the ECN during reciprocity suggests that these socially rewarding experiences may be encoded as more valued or salient than non-social rewards (Phan *et al.*, 2010; Fareri *et al.*, 2015).

We also found that the DMN exhibited enhanced connectivity with the superior frontal gyrus and superior parietal lobule—components of the frontoparietal control network—during reciprocity relative to violations of trust when interacting with close friends versus strangers. Activation within this network, involved in directing of control/attention to facilitate goal-directed behavior (Corbetta and Shulman, 2002; Vincent *et al.*, 2008), decodes tasks as a function of available performance incentives, which in turn facilitates behavior in a target detection task (Etzel *et al.*, 2016). Subnetworks within the frontoparietal system show connectivity with the DMN that appears to be specific to social and internally directed processes (Nihonsugi *et al.*, 2015; Xin and Lei, 2015; Bellucci *et al.*, 2019; Kam *et al.*, 2019). Therefore, interactions between the DMN and frontoparietal regions in our task suggest that the incentive value of reciprocity from a close friend may capture attention because of its overlap in self-other relevance based on past experience, which subsequently informs future predictions and choices (Barrett, 2017). An alternative, yet plausible account based on evidence that the DMN may represent personality information (Hassabis *et al.*, 2014), is that the DMN may be recruiting models of others’ personalities when interpreting outcomes of social interactions and relaying this information to attention and control related regions to facilitate future predictions and impressions of others.

The use of a network approach to task-based connectivity–nPPI–offers novel insight into the way in which canonical networks interact with other regions during social interactions involving trust. Previous work employing seed-based PPI has highlighted connectivity between the striatum and cognitive control regions (i.e., dACC, vlPFC) when deciding to trust outgroup members (Hughes *et al.*, 2016) and when experiencing violations of trust (Fouragnan *et al.*, 2013). While a seed-based approach can be informative regarding our understanding of communication between brain regions (Smith *et al.*, 2016), it is limited in its ability to provide broader insight to network dynamics during task-based conditions (Cole *et al.*, 2010). Evidence suggests a high degree of variability in connectivity of a region (e.g., precuneus) depending on the precise location of voxels within that region chosen as the ‘seed’ (Cole *et al.*, 2010). Second, specific regions or nodes within a known network may demonstrate heterogeneity in function and connectivity based on the nature and difficulty of the cognitive process being investigated: activation in overlapping components of the posterior cingulate cortex, for example, demonstrate functional connectivity with multiple neural networks (i.e., frontoparietal networks, cognitive control networks, DMN) as a function of in the moment cognitive and attentional demands (Leech *et al.*, 2011; Leech *et al.*, 2012). Related work has demonstrated that the precuneus exhibits increased connectivity with the DMN at rest, but increased connectivity with the frontoparietal network during task-states (Utevsky *et al.*, 2014). The treatment of entire networks as seeds in our approach allowed us to demonstrate that the DMN is critical for taking into account information about our social relationships when we are interacting with others and communicating that with components of frontoparietal networks possibly to indicate preferential orienting of attention. The ECN, on the other hand, appears less sensitive to the relative difference between social outcomes as a function of relationship closeness per se.

Our implementation of nPPI provides novel insight into the ways in which network connectivity is influenced by relationships during social interactions, yet the present study is not without limitations. First, while PPI analyses in general provide more in the way of characterizing neural interactions during psychological processes than connectivity analyses relying on simple correlation (Friston, 2009; Friston, 2011; Smith *et al.*, 2016), it does have multiple interpretations (Smith *et al.*, 2016). For example, clusters emerging as representing a significant PPI effect may reflect a change in connectivity due to the psychological context (e.g., reciprocity vs. violation of trust changes the interaction between the DMN and superior frontal gyrus). Alternatively, a significant PPI effect may indicate a change in the response of a target region to a specific context (e.g., reciprocity vs. violation) by activation in a seed (e.g., DMN). Second, these points also indicate that directionality of information flow can be ambiguous and is not directly indicated by a significant PPI result. Third, we chose to extract network time-series in this study using a 10-network parcellation as suggested by Smith and colleagues (Smith *et al.*, 2009); we chose this specific parcellation due to its more general anatomical definition of each network and lack of more targeted predictions about subnetworks or network nodes in our task. However, recent work (including studies cited here) has specified more fine-grained parcellations of the DMN and other networks (Dixon *et al.*, 2018; Buckner and DiNicola, 2019; Ji *et al.*, 2019). Future work looking at subnetwork connectivity during social interactions may be better able to characterize specific contributions to encoding of factors such as relationship closeness and outcome value, and may be able to identify how these processes may break down in samples with social difficulties. Fourth, we chose to keep rates of reciprocity in the trust game consistent across partners (50%), consistent with previous work from our group and to isolate any differences in neural responses to outcomes to partner context. However, it is possible that this may not best capture real-life dynamics within relationships. Additionally, while our results also appear specific to positive relative to negative social outcomes, other recent work (Park and Young, 2020) demonstrates a reduced role for one node within the DMN (rTPJ) when experiencing trust violations from a close friend that may be associated with a lower likelihood of updating impressions of close friends. Future work could probe changes in network connectivity during social reward processing and impression updating under more naturalistic contexts when violations of trust from friends and strangers may vary at different rates. Finally, while we are encouraged and intrigued by the findings reported here given that they survive rigorous correction for multiple comparisons (i.e., permutation testing), we interpret them cautiously, given our sample size.

Close relationships provide an important social context within which many of our day to day experiences occur and are characterized by repeated instances of trust and reciprocity. Building on previous findings indicating that social relationships influence the reward value of shared social experiences and that the DMN has a specialized role in social cognition, our results demonstrate that the context of close relationships shapes functional network dynamics during social interactions as a function of the closeness people feel towards a partner. These results and our approach have important implications for clinical samples characterized by difficulties with social cognition and forming relationships (e.g., autism, borderline personality disorder, schizophrenia) (Alcalá-López *et al.*, 2019) and individuals who may have a history of adverse social experiences (McLaughlin *et al.*, 2019). For example, hypoconnectivity has been observed between networks implicated in cognitive control and goal-directed processes (ECN) and processing of social stimuli (face-processing network) in relation to increased levels of autistic traits (Young *et al.*, 2015), while individuals with a history of early life caregiving adversity show hyperconnectivity at rest between the ventral striatum and mPFC that is associated with poor social regulation (Fareri *et al.*, 2017). Taking an nPPI approach to examine how relationship context and other social factors impact connectivity during social interactions may help more precisely pinpoint links between neural function and breakdowns in social behavior, subsequently leading to targets for intervention.

---

**Supplemental Figure 1.** Full correlations between DMN nodes and the DMN timecourse.

**Supplemental Figure 2.** Partial correlations between DMN nodes and the DMN.

