## Supplementary figures and images for "The influence of relationship closeness on default-mode network connectivity during social interactions"

### Supplemental Figure 1

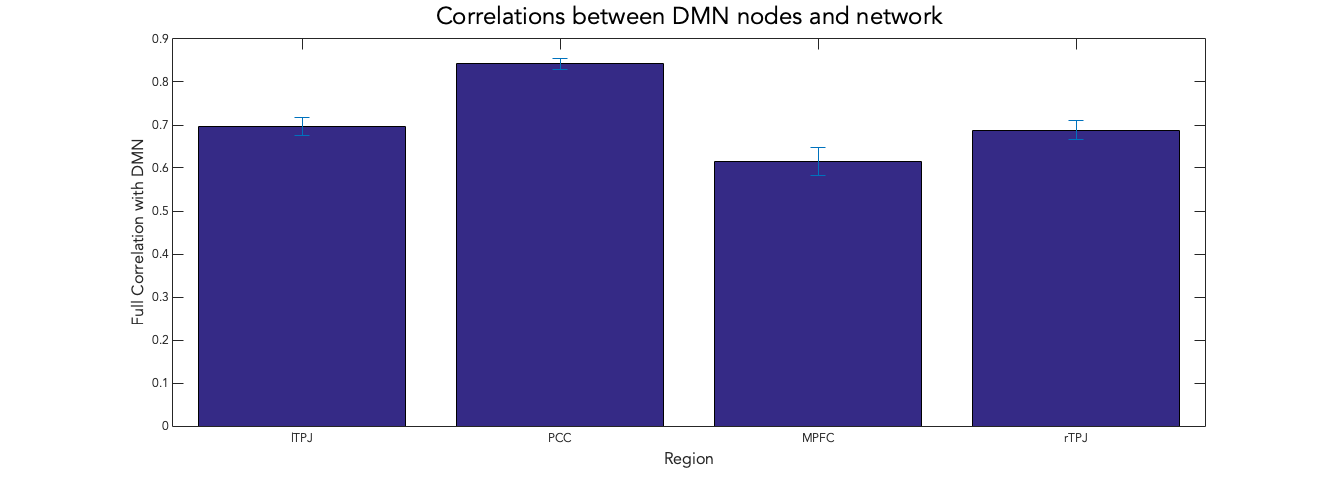

### Supplemental Figure 2

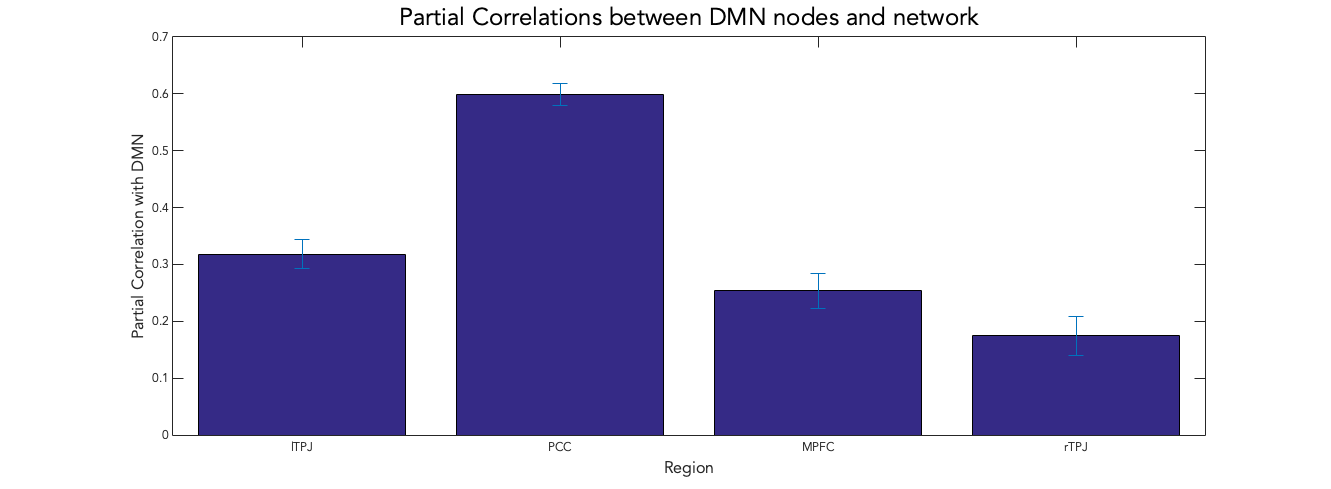
